# LPS-induced sepsis disrupts brain activity in a region- and vigilance-state specific manner

**DOI:** 10.1101/2024.12.25.630319

**Authors:** Susan Leemburg, Annu Kala, Athira Nataraj, Patricia Karkusova, Siddharth Baindur, Amritesh Suresh, Karel Blahna, Karel Jezek

**Affiliations:** Biomedical Center, Faculty of Medicine in Pilsen, Charles University, Alej Svobody 76, 323 00 Pilsen, Czech Republic; Neuroscience Research Center, Charité-Universitätsmedizin Berlin, Charitéplatz 1, D-10117 Berlin

**Keywords:** sepsis, sleep-wake architecture, state-space analysis, local sleep, aperiodic activity, oscillations

## Abstract

Sepsis-associated encephalopathy (SAE) is a common complication of sepsis and the systemic inflammatory response syndrome that leads to lasting consequences in survivors. It manifests as early EEG changes, that are region-, time- and state-specific, possibly reflecting distinct mechanisms of injury.

Here, we investigated the effects of 5mg/kg lipopolysaccharide (LPS) on hippocampal and cortical sleep-wake states, oscillatory and non-oscillatory neuronal activity, as well as on within and between state dynamics using state-space analysis.

LPS induced rapid-onset severe temporal and spatial vigilance state fragmentation, which preceded all other spectral changes by ∼90 minutes. Thereafter, LPS led to specific destabilization and increased delta oscillatory activity in wakefulness, but not NREM sleep, although state transitions remained largely normal. Instead, reduced NREM delta power resulted from aperiodic spectrum changes. LPS specifically reduced higher frequency hippocampal gamma oscillations (60-80Hz peak) in wakefulness, but not cortical high gamma or lower frequency gamma oscillations.

These results suggest that disruption of sleep-wake patterns could serve as an early indicator of sepsis and associated encephalopathy, independent of spectral changes. Moreover, treatment aimed at stabilizing vigilance states in early stages of sepsis might prove to be a novel option preventing the development of further pathological neurophysiology, as well as limiting inflammation-related brain damage.

## Introduction

Sepsis is a life-threatening complication of severe illness, consisting of a dysregulated systemic host response to an infection, accounting for 35% of hospital deaths.^1,2^ Up to 70% of sepsis patients suffer from sepsis-associated encephalopathy (SAE) occurring during early disease^3,4^ and 20-25% sepsis survivors experience post-sepsis syndrome^5^, which is associated with neurological symptoms such as sleep disturbances, impairment of cognitive and executive functions, and psychiatric conditions. Additionally, sepsis-related brain dysfunction manifests as delirium^4^, which independently predicts poor clinical outcomes^6,7^ and brain atrophy^8^.

While SAE may occur without clear neurological symptoms, EEG abnormalities such as EEG slowing, pathological theta waves, and triphasic waves have been found in sepsis patients^9–12^, although medical treatment and other conditions in the ICU hinder interpretation of these results somewhat. The severity of acute EEG abnormalities was directly related to mortality and disability, stressing the importance of early detection.^9,11,13^ Some EEG changes, including changes in theta and delta bands, remained in sepsis survivors, and were accompanied by hippocampal atrophy.^14^

Dysregulated sleep, fragmented by frequent awakenings, is one of the dominant neurologic signs in intensive care unit patients during acute sepsis.^4,15–18^ This fragmentation alone may limit the restorative functions of sleep, disrupt immune function and further predispose patients to SAE.^19–21^ As such, fragmented sleep is an important factor in the development on neurological symptoms in acute sepsis and in chronic post-sepsis syndrome^22^, where sleep fragmentation is associated with cognitive function, mental stability, and fatigue.^22–24^ Sleep-wake fragmentation is also present in rodent models of sepsis.^25,26^ We have previously shown that LPS not only affects oscillatory activity, but also the kinetics within hippocampal states in anaesthetized rats.^25^ We hypothesized that these facilitated the observed severe state fragmentation, forming a possible target for improving neurological outcomes in sepsis.

The varying EEG results recorded from patients suggest that SAE is heterogeneous, particularly at later disease stages, but also that different EEG phenomena could be present at different times. This also seems the case in animal models of sepsis and SAE: Cecal-ligation-and-puncture led to decreased NREM delta activity in acute sepsis^26^, while LPS injection caused increased wake delta in hippocampus, but not in prefrontal cortex.^27^ Wake theta was slowed in both hippocampus and prefrontal cortex, but only suppressed in cortex.^27^ Other studies in anaesthetized rats found no changes in delta, but described general slowing^28^ and reduced theta power instead^29^. As such, electrophysiological changes differ according to the used method of sepsis induction, time point of recording, vigilance state, and are at least in part region-specific.^26,27^ This may reflect different sources or mechanisms of damage, and thus different potential targets for intervention.

Here, we investigated the acute effects of high dose LPS on neural activity in cortex and hippocampus in the context of altered sleep-wake architecture in awake behaving animals, in order to better understand pathological within- and between-state dynamics, and their relation to other neurophysiological changes in acute sepsis.

## Methods

### Experimental setup

Hippocampal and cortical LFP and nuchal EMG were recorded on two days For 1 hour pre-injection and 6 hours post-injection. Rats were injected intraperitoneally with saline on the first recording day (BL) and with LPS on the second (5mg/kg in 0.9% NaCl). (Fig. 1A).

**Fig 1.**
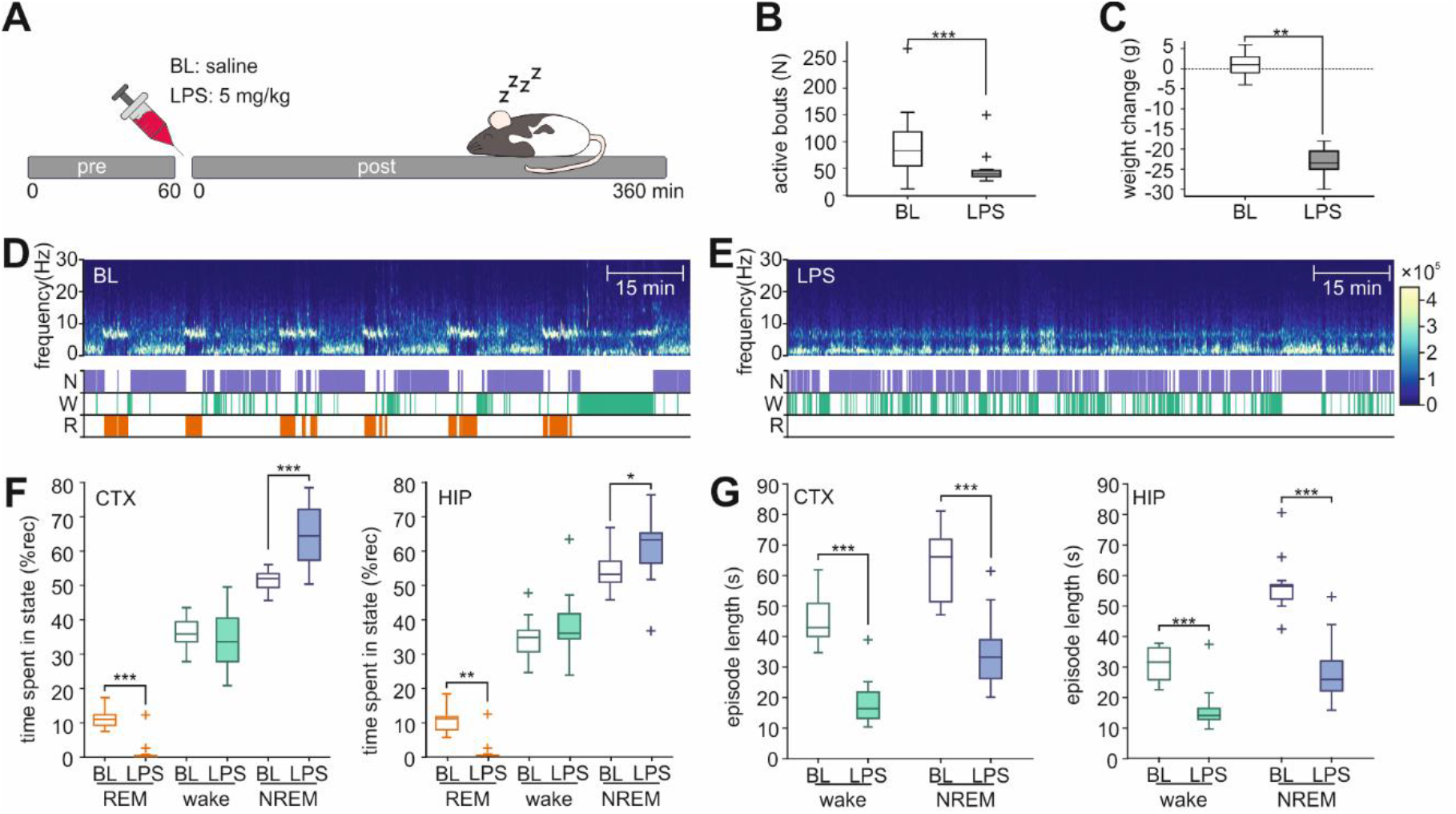
LPS causes sickness behavior and disrupts sleep-wake architecture. **A:** Schematic overview of the experiment. After a 1-hour pre-injection recording, rats were injected with saline or LPS, and recorded for another 6 hours. **B:** Active movement bouts post-injection at BL and after LPS. Box plots show median, 25th, and 75th percentiles. Values >1.5x interquartile range from the top or bottom of the box are displayed +; whiskers show minimum and maximum of the remaining values. **C:** LPS-induced weight loss 24 hours post-injection. **D+E:** Representative examples of spectrograms (top) and hypnograms (bottom), recorded 1 hour after saline injection (D) or after LPS injection (E). **F:** Time spent in wakefulness, NREM and REM sleep in cortex (CTX, left), and hippocampus (HIP, right). Brackets indicate significant post-hoc comparisons between BL and LPS per state, between-state comparisons are not shown. **G:** Episode lengths of wakefulness and NREM in cortex (CTX, left) and hippocampus (HIP, right). Brackets indicate significant post-hoc comparisons between BL and LPS per state, between-state comparisons are not shown. *p<0.05; **p<0.01; ***p<0.001

### Animals

13 male Long-Evans rats, aged 4 months at the time of recording, were individually housed in plexiglass cages with a 12h/12h light-dark cycle. During linear track training, rats were fed once daily in amounts that maintained a stable body weight. Water was available ad libitum. All experimental protocols were approved by the Ethical Committee of the Ministry of Education, Youth and Sports of the Czech Republic (approval no. MSMT-12084/2019) according to the Guide for the Care and Use of Laboratory Animals (Protection of Animals from Cruelty, Act No. 246/92, Czech Republic) and ARRIVE guidelines.

### Electrode implantation and recording

Rats were implanted with a 64-channel hyperdrive (Axona) containing 15 individually moveable tetrodes and one stereotrode constructed of twisted 17um-diameter platinum-iridium wire (California Fine Wire). The electrodes were coated with platinum to reduce the impedance to 120–200kΩ at 1kHz.

Briefly, after rats were anaesthetized using a mixture of ketamine and xylazine (100mg/kg and 10mg/kg, i.p., resp.) and 1-2% isoflurane in O_2_, a craniotomy was made over the right dorsal hippocampus (3.8 mm caudal, 3.2 mm lateral of bregma). After removal of the dura mater, the tetrode bundle was carefully lowered into the cortex to a depth of 1mm. Two Teflon-coated stainless steel wires were inserted into the neck muscles and served as EMG electrodes. Two stainless steel screws over the cerebellum served as reference. The craniotomy and electrodes were covered with paraffin wax, and the hyperdrive assembly was secured to the skull using stainless steel screws and dental acrylic. Rats were given carprofen (5mg/kg, s.c.) and Marboflaxin (5mg/kg, s.c.) for 3 days after surgery.

After recovery, individual tetrodes were lowered until they reached hippocampal CA1 over the course of ∼2 weeks. The stereotrode was left in its original location in the medial parietal association cortex (AP -3.6mm, ML 2mm, 1mm below brain surface). Electrode positions were histologically verified after the rats were sacrificed. All LFP and EMG signals were recorded at 24 kHz sampling rate using the computer-based Axona recording system (DacqUSB). Rat position and movement were tracked based on two infrared LEDs on the Axona headstage.

### Preprocessing

All LFP and EMG signals were downsampled to 6 kHz using Spikeinterface^30^ 0.100.6 in Python 3.9. Then, average LFP traces were calculated for each tetrode/stereotrode. Traces containing artifacts, as well as tetrodes located outside of CA1, were excluded from analysis. Overall, 7-15 hippocampal tetrodes were used for each rat. Cortical signals were excluded for one rat due to bad signal quality.

### Signal analysis

Sleep states were manually scored using AccuSleep^31^ in 4-second epochs. Scoring was performed independently for the cortical and hippocampal signals. For classification of the vigilance states, wake was characterized by low-amplitude, irregular LFP activity and highly variable EMG activity. NREM sleep, by contrast, was dominated by large amplitude slow waves and showed low EMG activity. REM sleep was defined by low amplitude LFP activity dominated by theta waves, with only occasional twitches in the EMG. The scorer was blinded to the experimental condition of the recording. Time spent per state, episode lengths, and state-divergence between cortex and hippocampus were analyzed. Scored epochs containing artifacts were excluded from power spectrum-and state-space analyses.

Power spectra were calculated using a sliding-window FFT analysis (4-s window, 2-s step) for each individual channel using Welch’s method in MATLAB 2019b (Hamming window, 50% overlap, 0.25-Hz resolution). Mean power spectra were then calculated per channel in time frames of interest for artifact-free epochs of each vigilance state. Coherence between cortical and hippocampal signals was calculated as magnitude-square coherence with a 0.25Hz resolution, and was then averaged for all hippocampal-cortical channel pairs to yield a coherence spectrum per rat. Periodic and aperiodic components of the power spectra were parametrized using FOOOF^32^. For each spectrum, the frequency range from 0.1 to 140 Hz was used with the following algorithm settings: peak width limits 0.5 and 12 Hz, maximum number of peaks: 6, minimum peak height: 0.05, and aperiodic mode: knee. Aperiodic spectral components were modelled using the following aperiodic fit AP(f) = 10^b^ × (1/(k+f^χ^)), where f is frequency, b is offset, k is the knee parameter, and χ is the spectrum slope. The knee parameter represents the bending point where the aperiodic fit transitions from horizontal to negatively sloped. Periodic components of the spectrum, representing putative oscillations, were modelled as Gaussian curves over and above the aperiodic background spectrum. Each oscillation has a center frequency (c), peak width (w), and center peak height (a), yielding the following for each detected frequency f: G(f)=a × exp(–(f – c)^2^/(2 × w^2^)).

Within- and between-state dynamics were analyzed using a 3-dimensional state space based on the sliding-window FFT time series, using three band-power ratios. Ratio 1 was calculated as 0.5-4Hz/6-10Hz, ratio 2 as 6-10Hz/100-200Hz, and ratio 3 as 10-15Hz/60-80Hz. The combination of frequency ranges was chosen based on approximate peak frequencies in power spectra, and on observed state separation in both cortex and hippocampus in pre-injection spectra. The resulting spectral-ratio-time series for all channels from a given brain region were then combined into a single series per region per rat using principal component analysis. The first principal component for each power-ratio was used to construct the state space. The time series were generally well represented by this component, which explained approximately 70% to 80% of variance at all time points for each ratio (for details, see Sup. S1). State-classification of points within state-space was based on manual scoring. Vigilance-state cluster positions in state space were calculated using median values for each power ratio. Within-state jitter was calculated as the Euclidian distance between two subsequent points, while state transition lengths were calculated as state-space path length of 7 state-space points centered on the state-change. Because of the relatively low number of pre-injection transitions, average transition length changes were bootstrapped by random selection of a number of post-injection transitions matching the pre-injection total. This was repeated 500 times to yield average transition lengths per rat.

Because tetrode positions changed between recording days, significantly altering signal characteristics, changes in power, coherence, and state space were normalized as percentage of pre-injection, and then averaged over all available channels per recorded area for each rat.

### Measures of sickness

To assess LPS-induced sickness severity, body weight and locomotor activity in the home cage were recorded. Active bouts were defined as periods where rats’ movement speed exceeded 2cm/s for at least 2 seconds. Home cage activity was analyzed independently from sleep-wake classification. Finally, disease severity was assessed by post-LPS mortality.

### Histology

After completion of the experiments, rats were overdosed with sodium pentobarbital (100mg/kg) and perfused transcardially with 0.01M phosphate-buffered saline (PBS) followed by 4% paraformaldehyde in PBS. Brains were cut in 50 µm thick coronal sections and Nissl stained. Electrode placement was then verified using a rat brain atlas^33^.

### Statistics

Statistical analyses were performed with JASP 0.18.1 and MATLAB 2019b. Changes in sleep state amounts and gamma oscillations were analyzed using repeated measures ANOVA (rmANOVA). Greenhouse-Geisser corrections for sphericity were applied where appropriate. Holm’s post-hoc test was used after significant factor interactions. For all other pairwise comparisons the non-parametric Wilcoxon signed-rank test (WSR) was used. Results where p<0.05 were considered statistically significant unless stated otherwise. Box plots in this paper show median and 25^th^ and 75^th^ percentiles. Values that are more than 1.5 times the interquartile range from the top or bottom of the box are displayed as a + sign; whiskers show the minimum and maximum of the remaining values.

## Results

### 1. Behavioral symptoms of the LPS-induced sepsis model

Rats’ home cage activity was relatively low in both BL and LPS recordings days, which took place during the light phase (Fig. 1A). After LPS injection, however, rats tended to remain inactive with a prone, flat posture, hardly any head scanning movements, half or fully closed eyes, and with some piloerection by the end of the recording period. At BL, rats were either in a normal, somewhat curled up, sleeping posture, or quietly sitting upright, when not actively exploring their cage. These behavior changes were also apparent in the lower number of activity bouts after LPS injection (WSR; z=2.691, p=0.008; Fig. 1B). Additionally, LPS injection led to significant weight loss 24 hours after injection, whereas saline injection did not (WSR; z=3.059, p=0.002; Fig. 1C).

### 2. LPS injection alters global and local sleep-wake architecture

LPS injection caused severe qualitative and quantitative disruptions of sleep-wake architecture in relation to states expression and in times spent in a given state, as well as in respective episode length (Fig. 1D-E). Effects were similar, but not identical in cortex (rmANOVA; day effect: F(1)=0.010, p=0.921; state effect: F(2)=144.922, p<0.001; day*state interaction: F(2)=7.652, p=0.002) and in hippocampus (rmANOVA; day effect: F(1)=0.060, p=0.808; state effect: F(2)=369.955,, p<0.001; day*state interaction: F(2)=12.283,, p<0.001). Qualitatively, we observed a significant, near-total reduction of REM after LPS injection in both brain regions, which was mostly compensated for with increased amounts of NREM (Fig 1F).

The remaining post-LPS wake and NREM were severely fragmented, with episode durations that were half as long as those recorded at baseline in cortex in both wake (WSR; z=2.417, p=0.013) and NREM (WSR; z=3.233, p=0.002). Episode lengths were similarly halved in hippocampus (wake: WSR; z=2.982, p=0.001; NREM: WSR; z=3.233, p=0.002) (Fig 1G). Combined analysis of all vigilance states, showed that, in addition to fragmentation, episode lengths became much more homogenous after LPS injection, with standard deviations per rat that were approximately half of BL (WSR; CTX_BL_ 104.06±4.34s vs. CTX_LPS_ 44.55±5.13s, z=3.059, p=0.005; HIP_BL_ 75.54±3.14s vs. HIP_LPS_ 35.00±4.95s, z=3.110, p=0.005; mean±s.e.m.).

Sleep state fragmentation occurred very rapidly after LPS injection (Fig. 2A). The number of state transitions was higher in hippocampus than in cortex before and after injection. LPS-induced state fragmentation was evident within the first 10 minutes after injection in both cortex and hippocampus (Fig. 2B). Although a further increase in fragmentation appeared to occur 60 minutes after injection, the number of state transitions at this time point was not significantly larger than at 10 minutes in either brain region (rmANOVA; CTX: recording time effect F(2)=4.478, p=0.017, day effect F(1)=13.794, p=0.001, time*day interaction F(2)=6.275, p=0.004. HIP: recording time effect: F(2)=7.959, p=0.001, day effect: F(1)=12.709, p=0.002, time*day interaction: F(2)=7.279, p=0.002). Fragmentation remained at baseline levels after saline injection.

**Fig 2.**
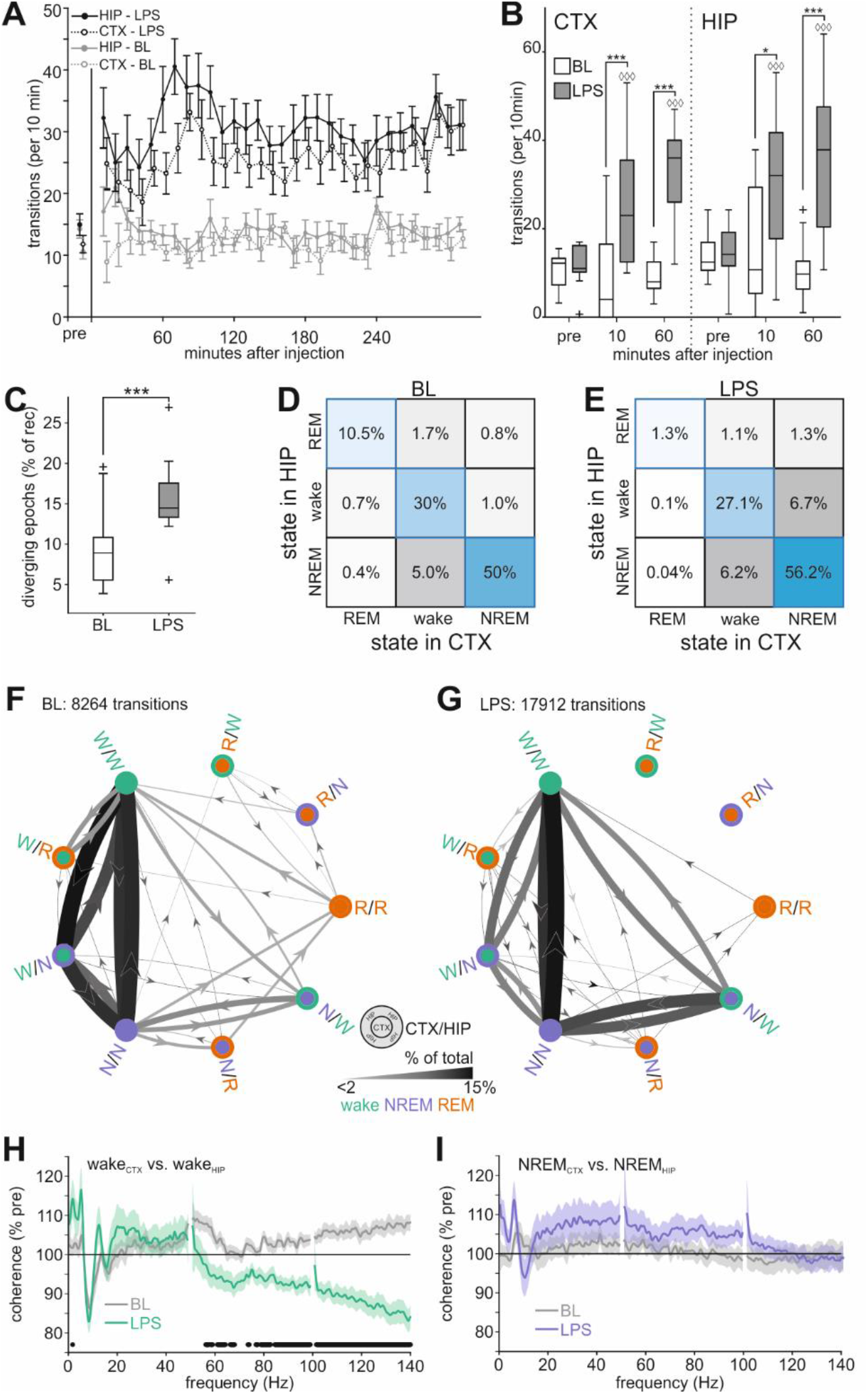
Rapid-onset sleep fragmentation and altered local sleep after LPS injection. **A:** Time course of the number of vigilance state transitions per 10-minute interval after injection in cortex (CTX) and hippocampus (HIP). Values are mean±s.e.m. **B:** State fragmentation pre-injection and 10 and 60-minutes post-injection. Brackets show differences between BL and LPS within a time point (*p<0.05; ***p<0.001). Diamonds indicate significant differences from same-day pre-injection (◊◊◊p<0.001). Box plots show median, 25^th^, and 75^th^ percentiles. Values >1.5x intequartile range from the top or bottom of the box are displayed as +; whiskers show minimum and maximum of the remaining values. **C:** Amount of state divergence in cortex and hippocampus. ***p<0.001. **D+E**: Matrix showing grouped hippocampal-cortical sleep-wake states at BL (D) and after LPS (E). **F+G**: Directed graphs showing transitions between cortex-hippocampus state combinations at BL (F) and after LPS (G). State-combinations are labelled as CTX/HIP and displayed as circles with the center representing the cortical and the edge representing the hippocampal state. Arrows show transition direction between state combinations. Arrow thickness and color is scaled according to the relative number of transitions. Transitions that occurred fewer than 20 times, and transitions between epochs of the same state-combination are not shown. **H:** Coherence between cortex and hippocampus when both are awake at BL (grey) and after LPS (green) as percentage of pre-injection. Values are mean±s.e.m. (N=12). Black dots below the plot show significant differences (paired t-test, p<0.01). **I:** Coherence between cortex and hippocampus when both are in NREM at BL (grey) and after LPS (violet) as percentage of pre-injection. Values are mean±s.e.m. (N=12).

However, even though overall effects of LPS on sleep-wake architecture were similar in cortex and hippocampus, the number of epochs where cortex and hippocampus were in different vigilance states was increased (WSR; z=-2.417, p=0.013, Fig. 2C), indicating that LPS may affect local sleep as well as global vigilance states. In baseline recordings, cortex and hippocampus spent up to 10% of recording time in diverging vigilance states; after LPS this increased to almost 15%. At BL, the most common diverging state combination was one where cortex displayed wake LFP patterns, while the hippocampus was in NREM sleep (Fig. 2D). In general, at baseline, the hippocampus entered NREM before the cortex, but also tended to transition back from NREM to wake later than the cortex, leading to a relatively high number of transitions between epochs with wake_CTX_-NREM_HIP_ and epochs where both regions were simultaneously in wake or NREM (NREM_CTX_-NREM_HIP_ and wake_CTX_-wake_HIP_) (Fig. 2F). After LPS injection, increased overall CTX-HIP state divergence manifested as higher numbers of epochs with wake_CTX_-NREM_HIP_, but also as a marked increase in NREM_CTX_-wake_HIP_ (Fig. 2E). This heightened state divergence was accompanied by altered state-transition patterns: transitions where hippocampus preceded cortex into NREM and followed into wakefulness remained common (wake_CTX_-NREM_HIP_ to NREM_CTX_-NREM_HIP_, or wake_CTX_-NREM_HIP_ to wake_CTX_-wake_HIP_), but the number of transitions between NREM_CTX_-NREM_HIP_ and NREM_CTX_-wake_HIP_ was notably increased. Additionally, transitions between the NREM_CTX_-wake_HIP_ and wake_CTX_-wake_HIP_ became much more common (Fig. 2G). That is, the normal cortical-hippocampal sequence of wake-NREM transitions was disrupted and both brain regions progressed through the sleep-wake cycle more independently of each other under LPS.

To investigate if the observed increased dissimilarity between cortex and hippocampus remained present even when both were in the same vigilance state, coherence spectra were analyzed for wake_CTX_-wake_HIP_ and NREM_CTX_-NREM_HIP_ epochs (Fig. 2H-I). In wakefulness, CTX-HIP coherence remained unaffected by LPS in lower frequencies, but showed a marked reduction in frequencies above 50Hz. By contrast, NREM coherence remained at BL levels at all investigated frequencies. So, LPS did not lead to overall changes in LFP similarity between brain regions in NREM, but led to reduced gamma coherence in wakefulness.

## 3. LPS destabilizes wakefulness, but not NREM

To further investigate LPS-induced changes in the dynamics of wakefulness and NREM, we employed a state-space consisting of three spectral power ratios (Fig. 3A-D; Ratio 1: 0.5-4Hz/6-10Hz, Ratio2: 6-10Hz/100-200Hz, Ratio 3: 10-15Hz/60-80Hz). This allowed us to investigate within- and between-state dynamics for cortex and hippocampus.

**Fig 3.**
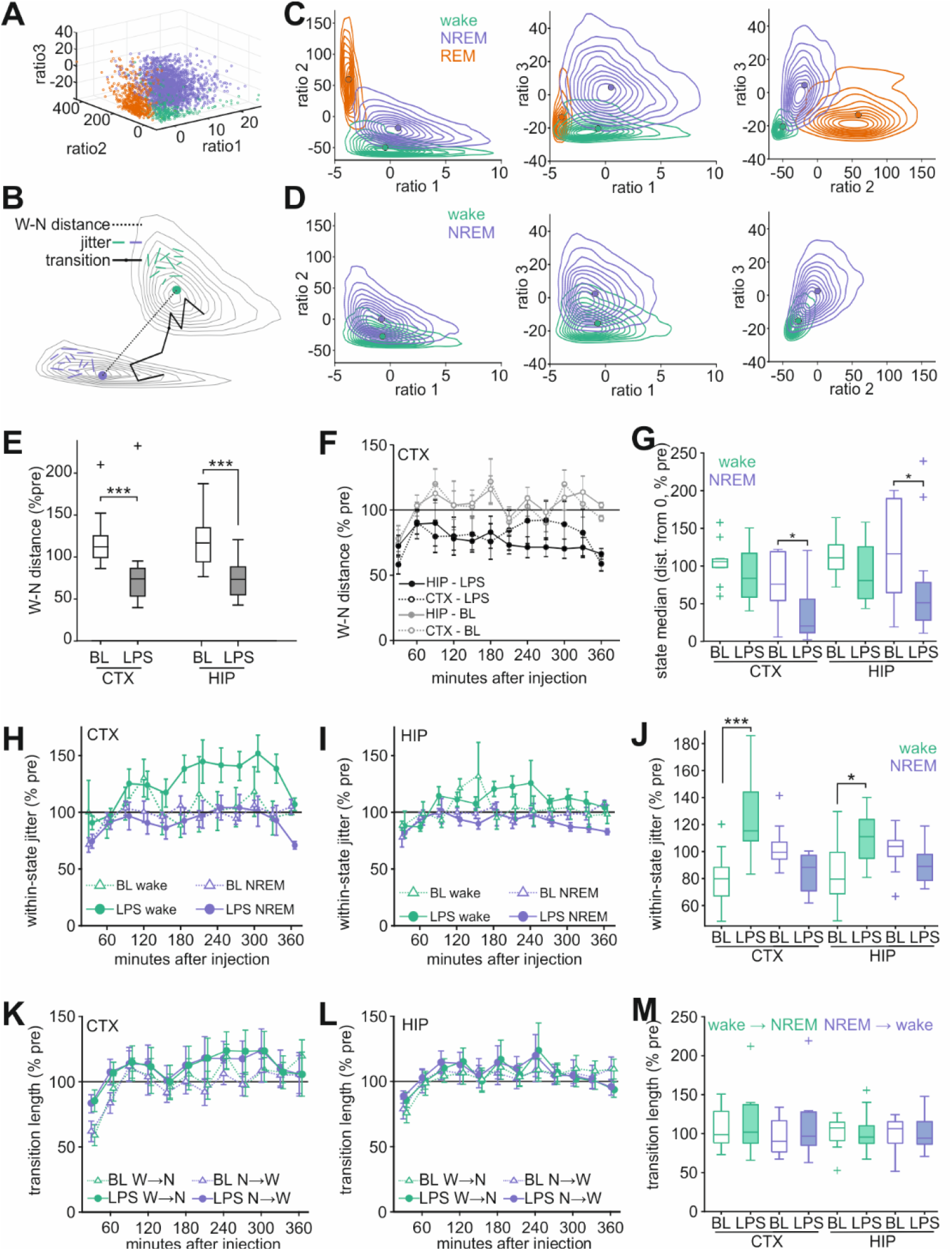
State-space analysis of LPS-related changes in LFP. **A:** Representative example showing separation of REM (orange), wake (green) and NREM (violet) after state-space decomposition along Ratio 1 (0.5-4Hz/6-10Hz), Ratio2 (6-10Hz/100-200Hz), and Ratio 3 (10-15Hz/60-80Hz). **B:** Schematic showing state distance, within-state jitter and transition length. Not to scale. **C+D**: Representative examples of post-injection state-space at BL (C) and after LPS (D) for all three spectral ratio combinations. Circles represent median values per state, with contours indicating samples per decile. **E:** Reduction of wake-NREM distances after LPS injection in cortex and hippocampus. ***p<0.001. **F**: Time course of wake-NREM distances in cortex and hippocampus. N=12 (CTX) and N=13 (HIP). **G:** Change in wake and NREM cluster medians at BL or after LPS. Values are shown as distance from origin, as percentage of pre-injection. **H+I**: Time course of within-state jitter in cortex (H) and hippocampus (I). **J:** Overall Within-state jitter at BL and after LPS. *p<0.05, ***p<0.001. **K+L:** Time course of transition lengths in cortex (K) and hippocampus (L) at BL and after LPS. Transitions from wake to NREM (green) and NREM to wake (violet) were analyzed. **M**: Overall transition lengths from wake to NREM and from NREM to wake were after LPS or saline injection. Box plots show median, 25^th^, and 75^th^ percentiles. Values >1.5x interquartile range from the top or bottom of the box are displayed as +; whiskers show minimum and maximum of the remaining values. Time course data shown as mean±s.e.m. N=12 (CTX) and N=13 (HIP).

The distance between wake and NREM clusters was significantly reduced after LPS injection in cortex and hippocampus (Fig. 3E; WSR; CTX: z=2.746, p=0.003; HIP: z=2.760, p=0.003), indicating increased spectral similarity between these states. Inter-state distances in LPS recordings were reduced compared to BL from approximately 60 minutes after injection, after which wake-NREM distances remained stable (Fig. 3F). Decreases in wake-NREM distance were mostly driven by changes in NREM, which resulted in cluster medians closer to the state space origin after LPS (Fig. 3G; WSR; CTX: z=2.118, p=0.034; HIP: z=2.132, p=0.033). Wake positions remained unaffected (WSR; CTX: z=0.471, p=0.677; HIP: z=1.825, p=0.068).

Within-state jitter, or the state-space distance between subsequent epochs, is a measure of state stability. More stable states, consisting of epochs with high similarity, occupy locations in close proximity within state space and show less jitter, whereas more instability is related to more jitter. LPS injection led to increased jitter in wake, but not NREM (Fig. 3H-J). This was the case in cortex (WSR; wake: z=-3.059, p=0.005; NREM: z=1.726, p=0.092) and in hippocampus (WSR; wake: z=-2.201, p=0.027; NREM: z=1.782, p=0.080) (Fig. 3J). Changes in jitter followed a similar time course to those in interstate distance, and became prominent after 60 minutes post-injection (Fig. 3H-I). This indicates a selective destabilization of wakefulness that occurred later than the onset of sleep-wake fragmentation, which arose within 10 minutes after LPS injection.

Although state transitions became much more numerous after LPS injection, and became less coordinated between cortex and hippocampus (Fig. 2A-G), the local spectral characteristics of transitions between wake and NREM within each brain region remained normal. Transition lengths, defined as distance covered in state space in a 12-second period prior to and after a state transition, were not significantly different for wake-to-NREM transitions (WSR; CTX: z=-1.098, p=0.301, HIP: z=0.105, p=0.946) and NREM-to-wake transitions (WSR; CTX: z=-1.569, p=0.129, HIP: z=-0.245, p=0.839).

## 4. LPS-related changes in oscillatory and aperiodic LFP activity

LPS injection resulted in widespread region- and state specific changes in the LFP power spectra, consisting of an initial broad-band increase in cortical power above ∼40Hz, followed by an overall loss of spectral power in both cortex and hippocampus that stabilized approximately 90 minutes after LPS injection (Fig. 4 A-D). Power spectra remained largely stable after saline injection (Sup. S2). Analysis of canonical spectrum bands confirmed an overall loss of power in wake and NREM in both cortex and hippocampus (Sup. S3-4), leading to reductions in sigma (10-12Hz), beta (12-30Hz) and gamma (30-90Hz) in all states and regions, as well as a loss of hippocampal wake theta, NREM theta in both cortex and hippocampus, and reduced hippocampal NREM delta.

**Fig 4.**
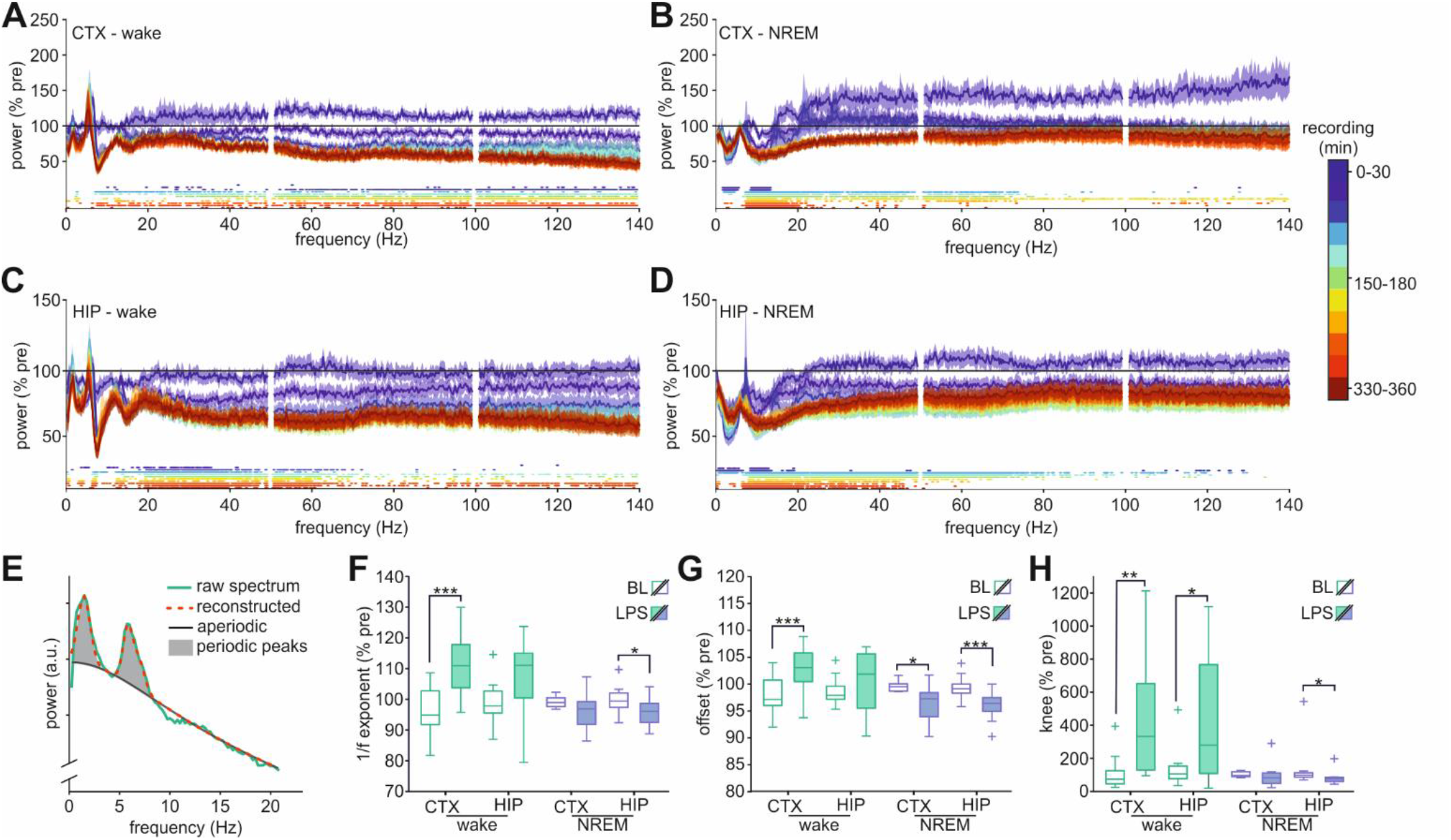
Effects of LPS injection on wake and NREM power spectra. **A-D:** Spectral power as percentage of pre-injection in cortex (A+B) and hippocampus (C+D) in wake (A+C) and NREM (B+D). Spectra are shown per 30-minute period after injection. Dots below the plots show significant differences compared to the same period at BL (paired t-test, p<0.01, also see Sup. S2). Values are mean±s.e.m. **E:** Example spectrum showing decomposition in aperiodic and periodic components. **F-H:** Changes in aperiodic spectrum parameters at BL (open) or LPS injection (shaded) in wake (green) and NREM (violet): 1/f exponent (F), offset (G), and knee (H). Box plots show median, 25^th^, and 75^th^ percentiles. Values >1.5x interquartile range from the top or bottom of the box are displayed as +; whiskers show minimum and maximum of the remaining values.

Loss of spectral band power may indicate loss of specific oscillations, but may also be related to non-oscillatory, aperiodic, effects. Since distinct mechanisms underlie such putative changes, separate analysis of oscillatory and aperiodic components of the power spectrum is warranted. LFP spectra recorded in the 90-360 minutes after injection were parametrized into their aperiodic and oscillatory components (Fig. 4E). Spectral parametrization resulted in reliable detection of four main oscillations: delta (1-4 Hz peaks), theta (6-9 Hz peaks) and two oscillations in the gamma range (γ1 at 20-40 Hz, γ2 at 60-80 Hz). Because of an apparent increase in higher frequency power in the first 30 minutes after LPS injection prior to the stabilization of power loss after 90 minutes, these two periods were analyzed separately for detected gamma peaks. In a subset of recordings, oscillations corresponding to spindles (10-12 Hz peaks) and beta (15-20 Hz peaks), and ripples (peaks >110Hz) were found. However, since they were only present in few rats and were absent at either a pre-or post-injection time point in most animals, these oscillations were not further analyzed.

### 4.1. Effects on aperiodic activity

Wake 1/f exponents were higher after LPS in cortex (WSR; z=-2.903, p=0.001, Fig. 4F), resulting in steeper spectrum slopes. Wake 1/f exponent changes did not reach statistical significance in hippocampus (WSR; z=-1.647, p=0.110). NREM exponents showed the opposite pattern, with significantly decreased exponents, resulting in shallower slopes in hippocampus (WSR; z=2.271, p=0.021), but not cortex (WSR; z=-1.689, p=0.102).

NREM spectrum offsets were reduced in both cortex (WSR; z=2.223, p=0.024) and hippocampus (WSR; z=3.040, p=0.007, Fig. 4G). However, wake offsets were increased in cortex (WSR; z=-2.197, p=0.027), but not hippocampus (WSR; z=-0.941, p=0.380).

Spectrum knee values showed a large and highly variable increase in wake in both cortex (WSR; z=-2.510, p=0.009, Fig. 4H) and hippocampus (WSR; z=-2.118, p=0.034). In NREM, knee values were reduced to ≈80% of pre-LPS, reaching statistical significance in hippocampus (WSR; z=2.062, p=0.040), but not cortex (WSR; z=1.156, p=0.278).

### 4.2. Effects on delta and theta oscillations

Detected baseline NREM delta peaks in cortex had peak frequencies of 1.72±0.05Hz, while hippocampal delta was slightly faster at 2.01±0.03Hz.

Hippocampal wake delta had a peak frequency of 1.41±0.05Hz. Delta peaks were slightly wider in hippocampus than in cortex (wake_HIP_: 1.71±0.08Hz, NREM_CTX_: 1.54±0.03Hz, NREM_HIP_: 1.90±0.05Hz).

Surprisingly, NREM delta oscillations remained largely unaffected by LPS (Fig. 5A-C). In both cortex and hippocampus, peak amplitudes (WSR; CTX: z=-1.511, p=0.147; HIP: z=-1.712, p=0.094), widths (WSR; CTX: z=1.867, p=0.067; HIP: z=1.293, p=0.216) and frequencies (WSR; CTX: z=1.600, p=0.123; HIP: z=1.363, p=0.191) were not significantly different at BL and after LPS, suggesting that observed reductions in delta band power (Sup. S3D) were not due to oscillatory changes. By contrast, LPS injection augmented delta oscillatory activity in wakefulness. Wake hippocampal delta amplitudes were increased after LPS (WSR; z=-2.666, p=0.004), but peak frequencies (WSR; z=0.889, p=0.426) and widths (WSR; z=0.889, p=0.426) remained at baseline levels. In cortex, no wake delta peaks were detectable at BL. After LPS, however, the number of rats with detectable oscillations showed a marked increase (frequency: 1.41±0.06Hz, width: 1.31±0.04Hz), likely resulting from delta amplitude increases as well (Fig. 5D).

**Fig 5.**
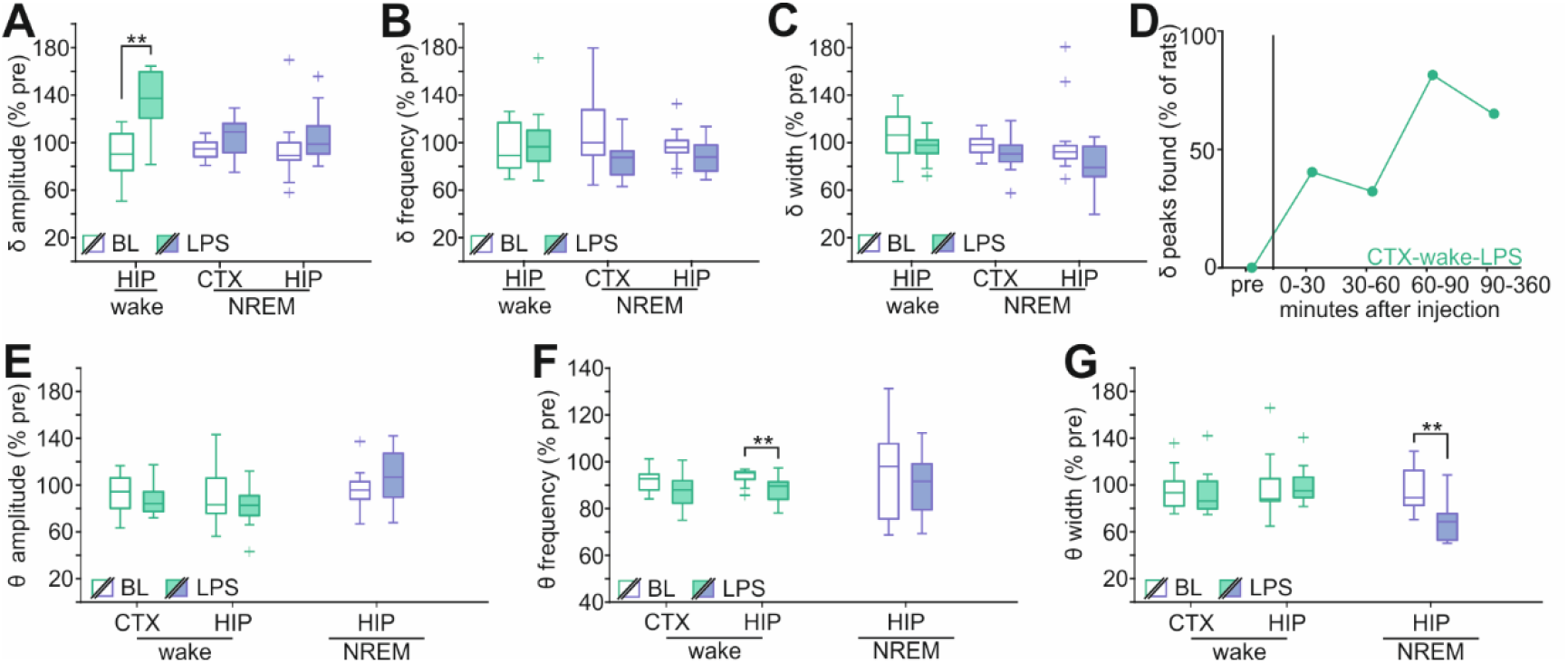
Effects of LPS on detected periodic delta and theta activity. **A-C:** Parameters of delta oscillations after saline (BL, open) or LPS injection (shaded) in wake (green) and NREM (violet). A: amplitude, B: frequency, C: width. Wake delta peaks were detected pre- and post-injection in 9/12 rats (HIP), NREM delta peaks in 11/2 rats (CTX) and 13/13 rats (HIP). **D**: Time course of detection of cortical wake delta oscillations after LPS. **E-G**: Parameters of theta oscillations at BL (open) or after LPS injection (shaded) in wake (green) and NREM (violet). E: amplitude, F: frequency, G: width. Wake theta peaks were detected pre- and post-injection in all rats (CTX: 12/12, HIP: 13/13). NREM theta was detected pre- and post-injection in 9/13 rats (HIP) and 4/12 rats (CTX). Box plots show median, 25^th^, and 75^th^ percentiles. Values >1.5x interquartile range from the top or bottom of the box are displayed as +; whiskers show minimum and maximum of the remaining values.

Theta oscillations in wakefulness were detected with baseline peak frequencies of about 7Hz (CTX: 7.12±0.05Hz, HIP: 7.04±0.04Hz) and wake peak widths of 1.90±0.04Hz (CTX) and 2.12±0.04Hz (HIP). NREM hippocampal theta peaks had similar peak frequencies (7.01±0.1Hz), but notably wider widths (4.07±0.31Hz).

Overall cortical wake theta oscillations remained mostly unaffected by LPS in cortex (WSR; amplitude: z=1.098, p=0.301, frequency: z=1.804, p=0.077, width: z=0.235, p=0.850). Hippocampal wake theta had slower peak frequencies after LPS (WSR; z=2.760, p=0.003), but no changes in amplitude (WSR; z=1.293, p=0.216) and width (z=-1.363, p=0.191) (Fig. 5E-G). Since hippocampal theta oscillations are strongly linked to locomotor activity, the observed effects on theta may solely be due to changes in behavior. However, when only epochs in quiet wakefulness (QW, movement speed <75^th^ percentile of post-LPS speed) were considered, LPS still caused a slowing of hippocampal theta, as well as a narrower theta peak (Sup. S5). QW theta amplitude was not significantly affected by LPS. QW theta peaks were not reliably detected in cortex.

NREM theta oscillations were reliably detected in hippocampus (9/13 rats), but not in cortex (4/12 rats). Hippocampal NREM theta oscillations became narrower after LPS injection (WSR; peak width: z=2.666, p=0.004), but theta frequency and amplitude were not affected (WSR; frequency: z=1.244, p=0.250; amplitude: z=-1.125, p=0.301).

### 4.3. Effects on cortical gamma oscillations

Two cortical oscillations were detected in the gamma band: a lower frequency γ1 peak and a higher frequency γ2 peak. These oscillations were readily detected in wakefulness (γ1 frequency: 38.6±0.9Hz, γ1 width: 10.7±0.9Hz; γ2 frequency: 61.1±0.9Hz, γ2 width: 9.1±0.6Hz), and in NREM (γ1 frequency: 34.2±1.1Hz, γ1 width: 10.1±0.7Hz; γ2 frequency: 66.0±1.2Hz, γ2 width: 10.2±0.8Hz).

Characteristics of the lower frequency γ1 oscillations remained unaltered by LPS in wakefulness, and γ1 remained detectable in most rats after injection (Sup. S6,7).

Wake γ2 amplitudes were reduced later in the recording compared to earlier in LPS-injected rats, but post-hoc tests showed no significant differences between LPS and BL at any time point (Fig. 6E; rmANOVA; recording time effect F(1)=16.100, p=0.007, day effect F(1)=0.046, p=0.833, time*day interaction F(1)=4.749, p=0.042). There were no changes in peak frequency (Fig. 6D; rmAVONA; recording time effect F(1)=0.899, p=0.355; day effect F(1)=0.004, p=0.985; time*day interaction F(1)=1.506, p=0.235) and width (Fig. 6E; rmANOVA; recording time effect F(1)=0.063, p=0.805, day effect F(1)=0.348, p=0.562, time*day interaction F(1)=0.241, p=0.629).

**Fig 6.**
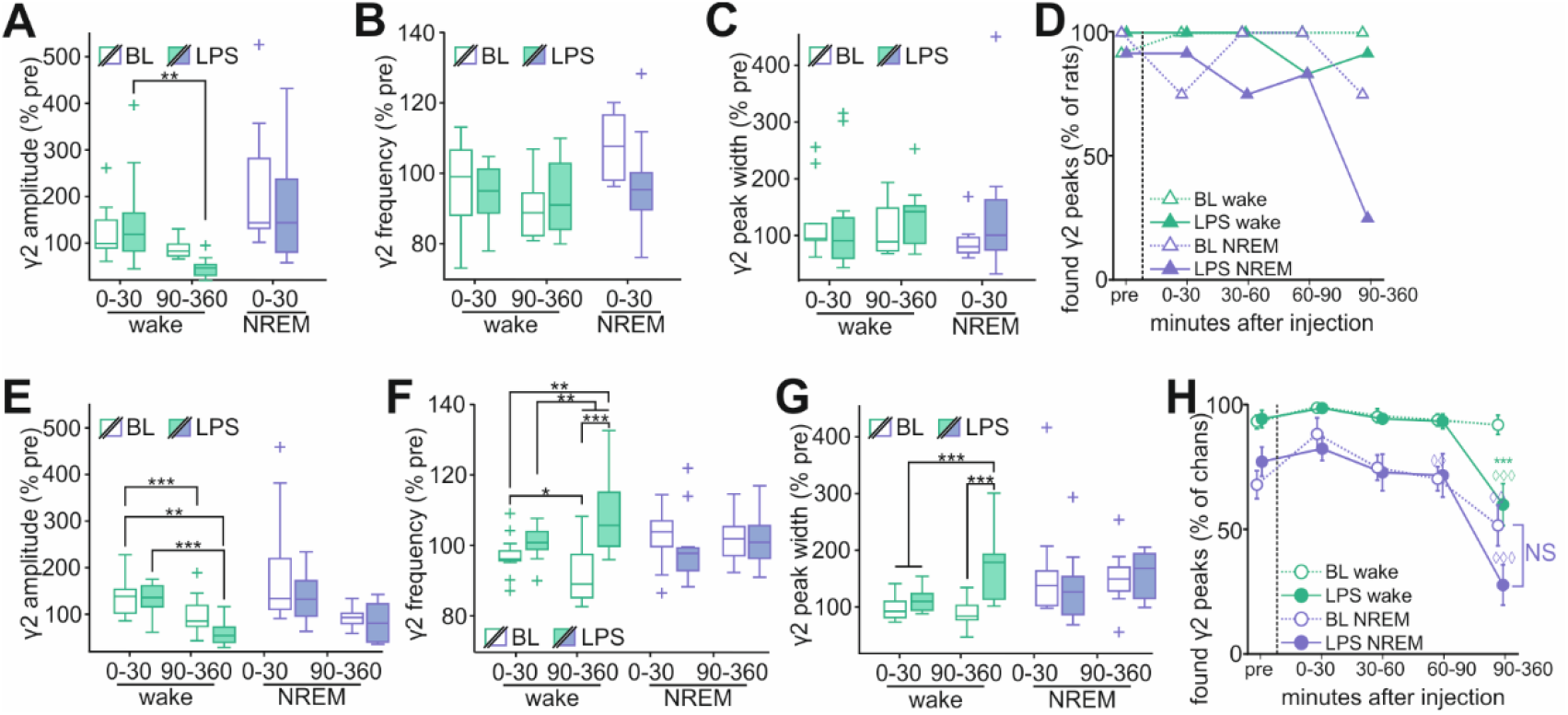
Effects of LPS on detected periodic gamma activity. **A-C**: Parameters of cortical high frequency gamma oscillations (γ2, 60-80Hz, detected pre- and post-injection in 11/12 rats (wake) and 8/12 rats (NREM) 0-30 and 90-360 minutes after injection. A: amplitude, B: frequency, C: width. **D:** Cortical γ2 peaks detection after LPS. **E-G:** Parameters of hippocampal high frequency gamma oscillations (γ2, 60-80Hz, detected pre- and post-injection in 13/13 rats). Peaks were analyzed 0-30 minutes and 90-360 minutes after injection. E: peak amplitude, F: frequency, G: width. **H:** Detected hippocampal γ2 peaks per channel. Box plots show median, 25^th^, and 75^th^ percentiles. Values >1.5x interquartile range from the top or bottom of the box are displayed as +; whiskers show minimum and maximum of the remaining values.

In NREM, both γ1 and γ2 became largely undetectable after 90 minutes and characteristics of either peak were not analyzed at this later time point (Fig. 6D,H; Sup S6,7). γ1 became undetectable both at BL and after LPS, whereas the loss of γ2 was more specific to LPS injection. In the first 30 minutes after LPS injection, γ1 was not significantly different from BL (Sup. S6). Cortical NREM γ2 was likewise unaffected by LPS (Fig. 6E-G; WSR; Amplitude: z=0.840; p=0.461; Frequency: z=1.680; p=0.109; Width: z=-0.140; p=0.945).

### 4.4. Effects on hippocampal gamma oscillations

Baseline hippocampal gamma peak characteristics were similar to those in recorded in cortex in wake (γ1 frequency: 36.4±0.6Hz, γ1 width: 10.3±0.5Hz; γ2 frequency: 61.0±0.8Hz, γ2 width: 9.6±0.4Hz), and in NREM (γ1 frequency: 35.9±0.5Hz, γ1 width: 9.8±0.4Hz; γ2 frequency: 63.2±1.2Hz, γ2 width: 11.0±0.5Hz).

Similar to cortex, hippocampal wake γ1 was not significantly affected by LPS injection in any of the analyzed parameters (Sup S6,8). NREM γ1 was likewise unchanged by LPS.

There was, however, a significant effect of time on detectability of hippocampal γ1 after injection at BL and after LPS. Fewer channels contained detectable oscillations after 90 minutes post-injection in both wake, although channels with detectable gamma remained present in all animals (Sup. S6,8). This reduction in detectability can be accounted for by the previously described time effects in γ1 amplitude.

In contrast to γ1, wake γ2 was significantly and specifically disrupted by LPS. γ2 amplitudes were significantly reduced after 90 minutes post-LPS, compared to BL and 0-30 minutes post-LPS (Fig. 6E; rmANOVA; recording time effect F(1)=94.396,, p<0.001, day effect F(1)=3.107, p=0.091, time*day interaction F(1)=5.250, p=0.031). γ2 peak frequencies were significantly higher after 90 minutes post-LPS, compared to a slight reduction at BL (Fig. 6F; rmANOVA; recording time effect F(1)=0.588, p=0.451, day effect F(1)=15.005, p=0.008, time*day interaction F(1)=17.379, p=0.004). Additionally, γ2 peaks became significantly wider after LPS injection (Fig. 6G; rmANOVA; recording time effect F(1)=8.424, p=0.008, day effect F(1)=21.235, p=0.001, time*day interaction F(1)=13.454, p=0.001). Detection of wake γ2 was reduced after LPS, but not at BL (Fig. 6H; rmANOVA; recording time effect F(1.772)=13.315,, p<0.001; day effect F(1)=4.655, p=0.041; time*day interaction F(1.772)=8.424, p=0.001).

Unlike wake γ2, NREM γ2 characteristics were not significantly affected by LPS injection. Peak frequencies (Fig. 6F; rmANOVA; recording time effect F(1)=4.589, p=0.047, day effect F(1)=1.389, p=0.255, time*day interaction F(1)=4.183, p=0.057) and widths (Fig. 6G; rmANOVA; recording time effect F(1)=0.846, p=0.370, day effect F(1)=0.337, p=0.569, time*day interaction F(1)=1.090, p=0.311) remained unaltered.

NREM γ2 amplitudes were not affected by LPS either, although a significant time effect was found (Fig. 6E; rmANOVA; recording time effect F(1)=11.968, p=0.003, day F(1)=1.239, effect p=0.281, time*day interaction F(1)=1.239, p=0.281). This time-dependent reduction in γ2 amplitudes paralleled a reduction in detectability of these oscillations in NREM. The number of channels with detected γ2 per rat was reduced after 90 minutes post-injection, but the difference between BL and LPS at this time point was not significant (Fig. 6H; rmANOVA; recording time effect F(4)=26.883,, p<0.001; day effect F(1)=0.295, p=0.592; time*day interaction F(4)=3.503, p=0.01).

## Discussion

Severe systemic inflammation, such as during sepsis, results in myriad alterations in brain activity and behavior. LPS injection resulted in severe sleep fragmentation, as well as a near complete suppression of REM sleep. While the sleep-wake fragmenting effects of inflammation are well documented^26,34,35^, the very rapid onset of these symptoms in sepsis has been described less frequently. Notably, effects on sleep-wake patterns preceded major changes in spectral power in both cortex and hippocampus by nearly 90 minutes in our study. A mouse LPS model showed that electrophysiological changes preceded neuroimmunological changes^36^. The rapid onset suggests a neural, rather than humoral, cause of state fragmentation. The later occurring spectral changes may instead be caused by slower humoral or glia-derived signals, be a consequence of state fragmentation itself, or an interaction of both. Sleep fragmentation by itself exacerbates febrile responses to LPS^37^, disrupts the blood-brain-barrier^38^, and leads to increased mortality after sepsis^39^, even when overall sleep and wake times remained unaffected. Since the observed time course in spectral changes coincided with the appearance of sickness behaviors in mice^36^ and seems linked to cognitive recovery^40^, prevention of state-specific EEG abnormalities might promote better outcomes after sepsis. Stabilization of sleep-wake states might prevent downstream spectral abnormalities, but also mitigate direct effects of sleep disruption on inflammatory responses.

In addition to temporal state fragmentation, sleep-wake patterns became more spatially fragmented: cortex and hippocampus showed increased diverging local sleep and wake activity after LPS. Sleep is regulated by both global, bottom-up processes^41,42^ and by local processes that affect brain regions^43–46^, local neuronal populations^47,48^, or even individual neurons^49^. This combination of local and global vigilance state regulation results in periods where different brain areas express different sleep state signatures within the same brain, sometimes for prolonged periods of time^44,50,51^. Sleep-state divergence occurs particularly around transitions in global sleep states, but can also occur outside of transitional states^50^. Here, we found increased state-divergence in acute systemic inflammation. This may be due to a higher overall number of transitions, but LPS-related changes in transition patterns also pointed to disruption of the normal cortico-hippocampal state-transition sequences, possibly indicating a weakening of global sleep-wake regulation processes^52–54^.

State-space analysis showed increased similarity between wake and NREM under LPS, an effect described previously in anaesthetized rats^25^. Increased state similarity was also reflected in aperiodic spectrum changes, which showed generally opposing patterns of change in wake and sleep in cortex and hippocampus. In addition, wake-specific destabilization was present in both regions. We previously hypothesized that high state similarity, in combination with heightened within-state instability, could facilitate sleep-wake fragmentation.^25^ However, the large delay between the onset of fragmentation and changes in inter-state similarity and jitter precludes such a causal role. Interestingly, state fragmentation and changes in power spectrum appeared much later after LPS injection under urethane in rats^25^ and mice^36^, than in our awake behaving rats.

Effects of LPS on intracortical and intrahippocampal activity were region-specific and vigilance-state-specific. Surprisingly, we did not detect any changes in delta oscillatory activity in either cortex or hippocampus in NREM sleep. Previous studies have shown a loss of NREM delta band power in sepsis^26^, which we replicate here.

However, the overall decrease of NREM delta power found in the current paper resulted from aperiodic changes in spectrum slope and offset, rather than from reduced delta oscillations. By contrast, wake delta oscillation amplitudes in both cortex and hippocampus were increased by LPS. Both delta activity^55,56^ and spectrum slopes^57^ are markers of sleep homeostasis under healthy conditions, but their underlying mechanisms in sepsis remain to be elucidated. Hippocampal wake theta oscillations were slowed after LPS, matching previous results.^27^ However, even when movement speeds were matched in BL and LPS groups, wake theta was slower after LPS than at BL, suggesting that this change was not solely the result of altered locomotor activity. These changes in lower frequency oscillations point to a less active wakefulness state^58,59^, which matched observed behaviors. They are also consistent with the spectral slowing observed in sepsis patients.^9,11–13^

LPS-related suppression of high frequency neural activity has gained attention as a possible cause of inflammation-related delirium and other cognitive symptoms. Delirium is a common complication during sepsis, and a strong predictor for increased mortality and poor cognitive outcomes.^6,7,60^ Indeed, loss of high frequency brain activity, i.e. beta power, was specifically related to delirium in sepsis patients^61^. Reductions in gamma power have been shown in both rat and mouse LPS models of sepsis^36,62^, although increases have been described in the cecal-slurry model.^40^ Moreover, evoked and non-evoked gamma are differentially suppressed after LPS.^62^ Here, we show distinct effects of LPS on slower and faster oscillations within the gamma band, mostly in wakefulness. Hippocampal and cortical lower frequency gamma oscillations were not specifically affected by LPS, but showed significant time-related reductions in amplitude. Higher frequency gamma oscillations were specifically weakened in wakefulness in both cortex and hippocampus after LPS, but not after saline injection. This reduction in regional gamma oscillations was accompanied by reduced cortico-hippocampal coherence in gamma frequencies. Additionally, our results show that experiments regarding the role of gamma oscillations in LPS-related symptoms, such as delirium, requires careful experimental design to account for sepsis-or treatment-related effects as well as possible confounders like time and vigilance state. Moreover, gamma oscillations should not be analyzed as a single broad-frequency band, as this may obscure disease-relevant relevant changes.

Aperiodic spectrum knees values showed large increases in wakefulness after LPS. Spectrum knee values can reflect intrinsic neuronal timescales, where higher knee values indicate faster time constants of a neuronal network, or a shorter time over which that network integrates and updates information.^63^ In humans, longer prefrontal cortical timescales predict working memory performance, while they shorten with age.^63^ Inflammation-induced compression of timescales limits the ability of hippocampal and cortical networks to integrate inputs sufficiently long to generate appropriate and sufficiently stable output patterns. This may lead to further discoordination between brain regions, and worsen cognitive symptoms in sepsis.

The overall sepsis-induced changes in wake LFP oscillations, timescale characteristics, and within-state-stability, combined with increased spatial fragmentation, points to a brain that is unable to maintain activity- and functional connectivity patterns required for cognition and execution of stable cognition and behavior. Clinically, such a neuropathological state would manifest as delirium or more severe neurological symptoms.^4^ Disruption of sleep-wake patterns could serve as an early indicator of sepsis and associated brain dysfunction, like SAE. Moreover, if the spectral changes observed in the current paper not only follow state fragmentation, but are a direct or indirect consequence, treatment aimed at stabilizing and restoring normal sleep-wake patterns in early stages of sepsis might prove to be a novel option in treating SAE and preventing more severe sepsis-related brain damage.

## Abbreviations

rmANOVA: repeated measures analysis of variance;
CTX: cortex;
HIP: hippocampus;
BL: baseline;
LPS: lipopolysaccharide;
LFP: local field potential;
EMG: electromyogram;
WSR: Wilcoxon signed-rank test;
NREM: non rapid eye movement sleep;
REM: rapid eye movement sleep

## Acknowledgements

This work was supported by Charles University Projects START/MED/107 and Cooperatio/NEUR, by project CZ.02.1.01/0.0/0.0/16_019/0000787 “Fighting INfectious Diseases,” awarded by the MEYS CR and financed from EFRR, and by project nr. LX22NPO5107 (MEYS): Financed by EU - Next Generation EU.

