## supplement for "LPS-induced sepsis disrupts brain activity in a region- and vigilance-state specific manner"

### Sup. S1

| day | ratio 1 | ratio 2 | ratio 3 |
|---|---|---|---|
| BL | 67.4±4.5 | 79.0±3.7 | 82.8±1.8 |
| LPS | 65.8±3.8 | 82.8±3.2 | 78.7±2.9 |

**Sup. S1:** Variance explained by first principal component per ratio at BL and after LPS. Values shown are mean±s.e.m.

### Sup. S2

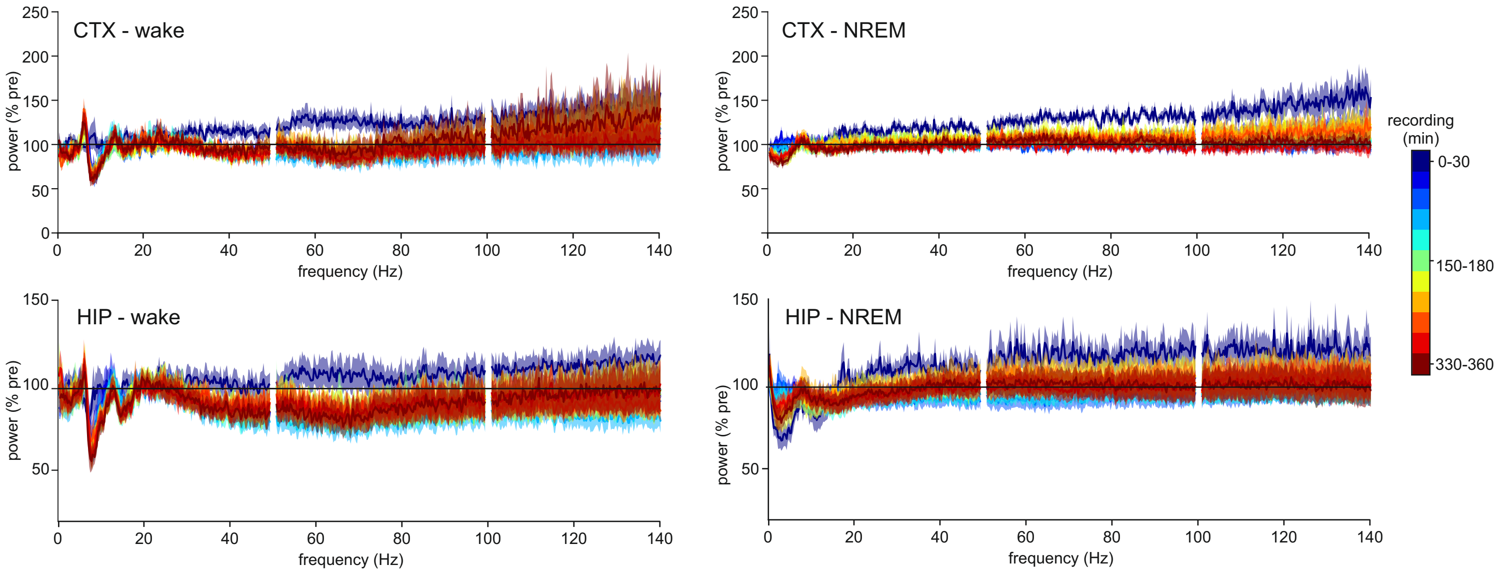


**Sup. S2:** Power spectra in saline injected rats
**A-D**: Spectral power as percentage of pre-injection in cortex (A+B) and hippocampus (C+D) in wake (A+C) and NREM (B+D). Spectra are shown for each 30-minute period after injection.

#### Sup. S3

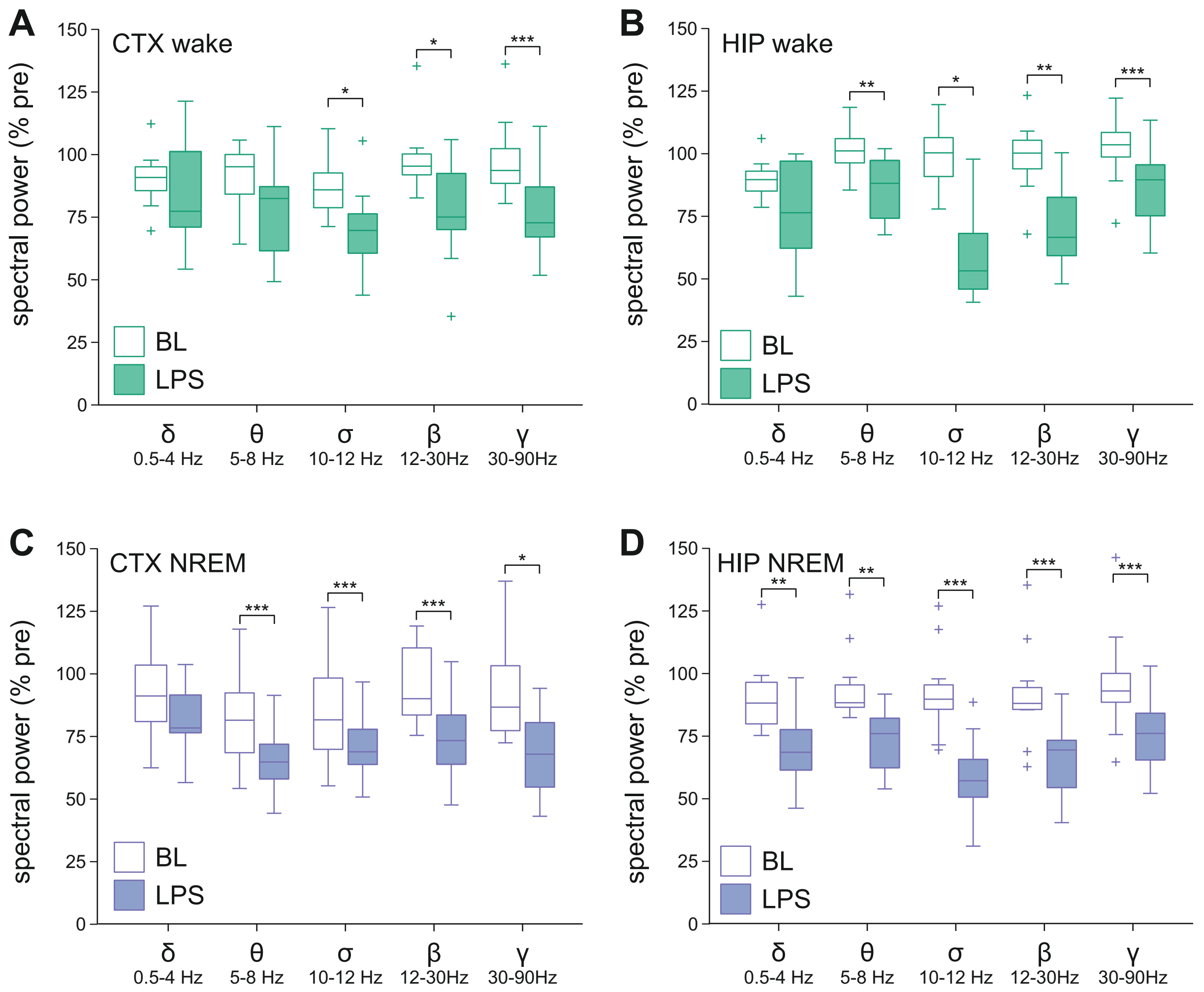


**Sup. S3:** Power per band after saline and LPS injection.
**A-D:** Changes in spectral power in canonical bands at baseline or after LPS injection in CTX (A,C) and HIP (B,D) for wakefulness (A-B) and NREM sleep (C-D). *p<0.05, **p<0.01, **p≤0.001. Differences per band were analyzed for each brain region and vigilance state using Wilcoxon signed-rank test (WSR). Test statistics and p-values are shown in table S3.

#### Sup. S4

| | | Wake | | NREM | |
|---|---|---|---|---|---|
| | Band | WSR | p | WSR | p |
| CTX | Delta | 0.392 | 0.733 | 1.726 | 0.092 |
| CTX | Theta | 1.490 | 0.151 | 2.903 | 0.001*** |
| CTX | Sigma | 2.432 | 0.012* | 3.059 | <0.001*** |
| CTX | Beta | 2.353 | 0.016* | 2.981 | <0.001*** |
| CTX | Gamma | 3.059 | <0.001*** | 2.275 | 0.021* |
| HIP | Delta | 1.363 | 0.191 | 2.551 | 0.008** |
| HIP | Theta | 2.551 | 0.008** | 2.900 | 0.002** |
| HIP | Sigma | 2.201 | 0.027* | 2.970 | 0.001*** |
| HIP | Beta | 2.830 | 0.002** | 3.040 | <0.001*** |
| HIP | Gamma | 2.970 | 0.001*** | 2.621 | 0.006** |

**Sup. S4:** Statistics for data shown in Fig. S2.
Test statistics and p-values for comparisons of baseline and post-LPS spectrum band power.

### Sup. S5

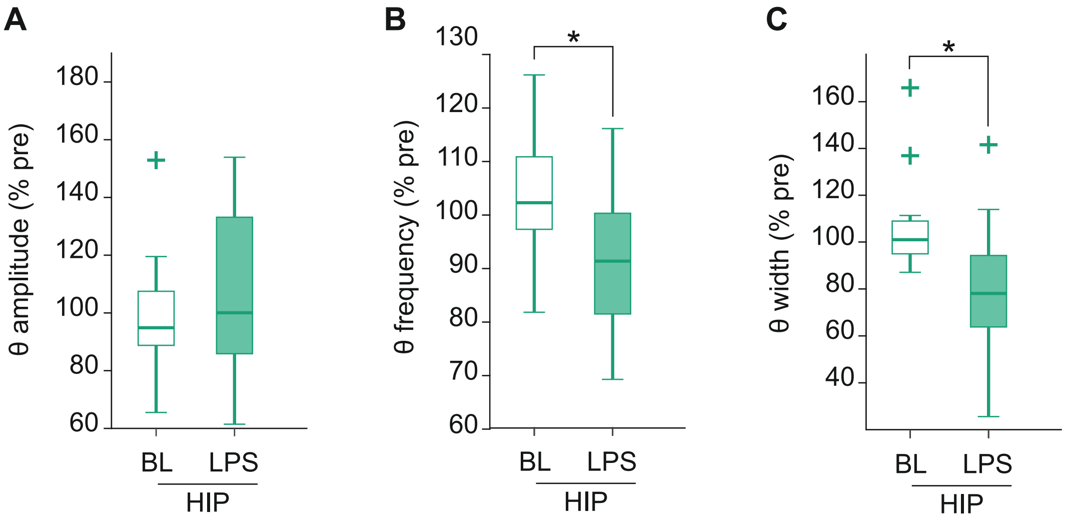


**Sup. S5:** Hippocampal theta oscillations during quiet wakefulness.
Peak parameters of detected theta during quiet wakefulness (QW) in HIP at baseline and after LPS injection (**A**: peak amplitude, **B**: peak frequency, **C**: peak width).
Epochs where movement speed was below the 75th percentile of in movement speed after LPS injection were selected as QW. For each rat, pre- and post-injection epochs matching this criterium were selected pre- and post-injection at BL and after LPS. While QW theta amplitude was not significantly affected (WSR, z=-0.804, p = 0.455), LPS injection led to narrower theta peaks in QW (WSR, z=2.411, p = 0.013) with lower peak frequencies (WSR, z=2.411, p = 0.013). QW theta peaks were not reliably detected in cortex. Theta peaks were present in 7/12 rats at BL and after LPS, but only 4 of these rats had a detectable peak at both the pre- and post-injection time point on each day.

#### Sup. S6

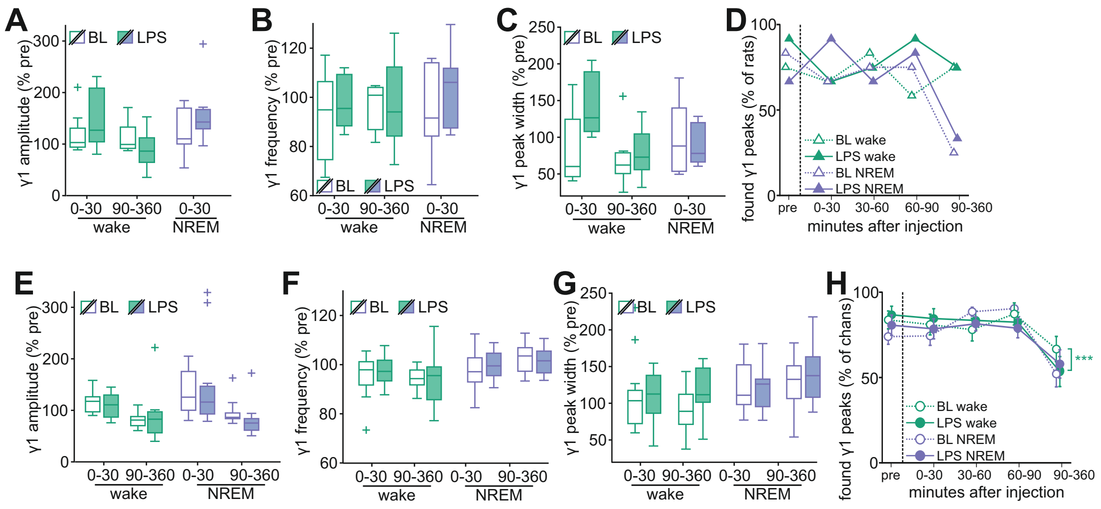


**Sup. S6:** Effects of LPS on detected periodic gamma activity.
**A-C**: Parameters of cortical low frequency gamma oscillations (γ1, 25-45Hz, detected pre- and post-injection in 7/12 rats) 0- 30 and 90-360 minutes after injection. A: amplitude, B: frequency, C: width. **D**: Cortical γ1 peak detection after LPS.
**E-G**: Parameters of hippocampal low frequency gamma oscillations (γ1, 25-45Hz, detected pre- and post-injection in 13/13 rats) 0-30 minutes and 90-360 minutes after injection. E: amplitude, F: frequency,G: width. **H:** Detected hippocampal γ1 peaks per channel.
Box plots show median, 25th, and 75th percentiles. Values >1.5x interquartile range from the top or bottom of the box are displayed as +; whiskers show minimum and maximum of the remaining values.

#### Sup. S7

| | CTX γ1 | Wake | | NREM | |
|---|---|---|---|---|---|
| | | rmAVONA | p | WSR | p |
| amplitude | recording time effect | 10.199 | 0.033 | -0.674 | 0.625 |
| | day effect | 0.516 | 0.512 | | |
| | time*day interaction | 0.505 | 0.516 | | |
| frequency | recording time effect | 0.818 | 0.389 | -2.023 | 0.063 |
| | day effect | 0.818 | 0.389 | | |
| | time*day interaction | 0.818 | 0.389 | | |
| width | recording time effect | 0.979 | 0.378 | 0.944 | 0.438 |
| | day effect | 0.950 | 0.385 | | |
| | time*day interaction | 0.608 | 0.479 | | |

**Sup S7:** Statistics for data shown in Fig. S6 A-C.
Test statistics and p-values for comparisons of baseline and post-LPS cortical γ1 oscillations.
Results for Wake are rmANOVA with factors recording time and day; results for NREM are WSR.

#### Sup. S8

| | HIP γ1 | Wake | | NREM | |
|---|---|---|---|---|---|
| | | rmAVONA | p | rmANOVA | p |
| amplitude | recording time effect | 14.860 | <0.001 | 24.00 | <0.001 |
| | day effect | 0.027 | 0.871 | 0.045 | 0.834 |
| | time*day interaction | 0.412 | 0.527 | 0.951 | 0.340 |
| frequency | recording time effect | 2.929 | 0.100 | 1.935 | 0.178 |
| | day effect | 0.004 | 0.948 | 0.015 | 0.904 |
| | time*day interaction | 0.055 | 0.817 | 0.920 | 0.348 |
| width | recording time effect | 0.577 | 0.455 | 3.072 | 0.094 |
| | day effect | 0.852 | 0.366 | 1.015 | 0.325 |
| | time*day interaction | 2.022 | 0.168 | 1.251 | 0.275 |
| found peaks | recording time effect | 5.623 | 0.001 | 20.712 | <0.001 |
| | day effect | 0.330 | 0.571 | 0.129 | 0.723 |
| | time*day interaction | 0.501 | 0.682 | 2.227 | 0.098 |

**Sup S8:** Statistics for data shown in Fig. S6 E-H.
Test statistics and p-values for comparisons of baseline and post-LPS cortical γ1 oscillations.
Results for for wake and NREM are rmANOVA with factors recording time and day. P-values for found peaks are Greenhouse-Geisser adjusted.
